# Circular RNA aptamers ameliorate AD-relevant phenotypes by targeting PKR

**DOI:** 10.1101/2024.03.27.583257

**Authors:** Xin Feng, Bo-Wen Jiang, Si-Nan Zhai, Chu-Xiao Liu, Hao Wu, Bang-Qi Zhu, Meng-Yuan Wei, Jia Wei, Li Yang, Ling-Ling Chen

## Abstract

Here, we delineated the remarkably elevated neuroinflammation accompanied by progressive activation of double-stranded RNA (dsRNA)-activated Protein Kinase R (PKR) and PKR-related dsRNA pathways in hippocampus of 5×FAD mice upon Alzheimer’s disease (AD) progression. AAV-delivery of circular RNAs possessing short-imperfect duplex regions (ds-cRNAs) to neurons and microglia effectively dampened excessive PKR activity with little toxicity, accompanying with reduced neuroinflammation and amyloid-beta (Aβ) plaques, resulting in neuroprotection and enhanced capability of spatial learning and memory in AD mouse models. These findings suggest a therapeutic potential of ds-cRNA aptamers as PKR inhibitors in AD therapy.

## Main

Alzheimer’s disease (AD) is an age-related neurodegenerative disease characterized by progressive memory decline and cognitive dysfunction, marked by deposition of amyloid-beta (Aβ) plaques and neuroinflammation^1, 2^. Neuroinflammation plays a central role in its etiopathogenesis^3, 4^, owing to exacerbate Aβ and Tau pathologies^5^. Double-stranded RNA (dsRNA)-activated Protein kinase R (PKR) participates in key inflammatory pathways with its activated phosphorylated form^6^. Depleting PKR has shown promise in rescuing synaptic and learning deficits in AD mice^7–10^, making it a potential therapeutic target. However, current small molecule PKR inhibitors often lead to side effects such as organ toxicities, hindering their clinical application^11^. Notably, circular RNAs with 16-26bp imperfect RNA duplexes (ds-cRNAs) exhibit potent PKR inhibition^12^, suggesting their potential in addressing excessive PKR activation in AD.

Consistent with the known PKR accumulation and p-PKR activation in AD brains^13–15^, we noted a gradual increase in PKR and phosphorylated PKR (p-PKR) levels in the hippocampus of 5×FAD mice, an AD model with accelerated amyloid deposition by expressing human APP and PSEN1 transgenes with a total of five related mutations. The elevation in PKR and p-PKR levels starts at 6 months of age, contrasting with the levels observed in wild-type (WT) controls (Fig.1a and Extended Fig.1a). Additionally, pro-inflammatory cytokines^15–18^ exhibited heightened expression in AD mouse hippocampus also starting from 6 months of age (Extended Data Fig.1b) and became much higher than WT controls at 10 months (Fig. 1b). Significantly, RNase L demonstrated a progressive increase consistent with pro-inflammatory cytokines, with higher activities observed in 10-month-old 5×FAD mice compared to WT controls (Fig. 1c and Extended Fig. 1c), indicating the involvement of circRNAs in AD-related neuroinflammation^12^.

**Fig. 1.**
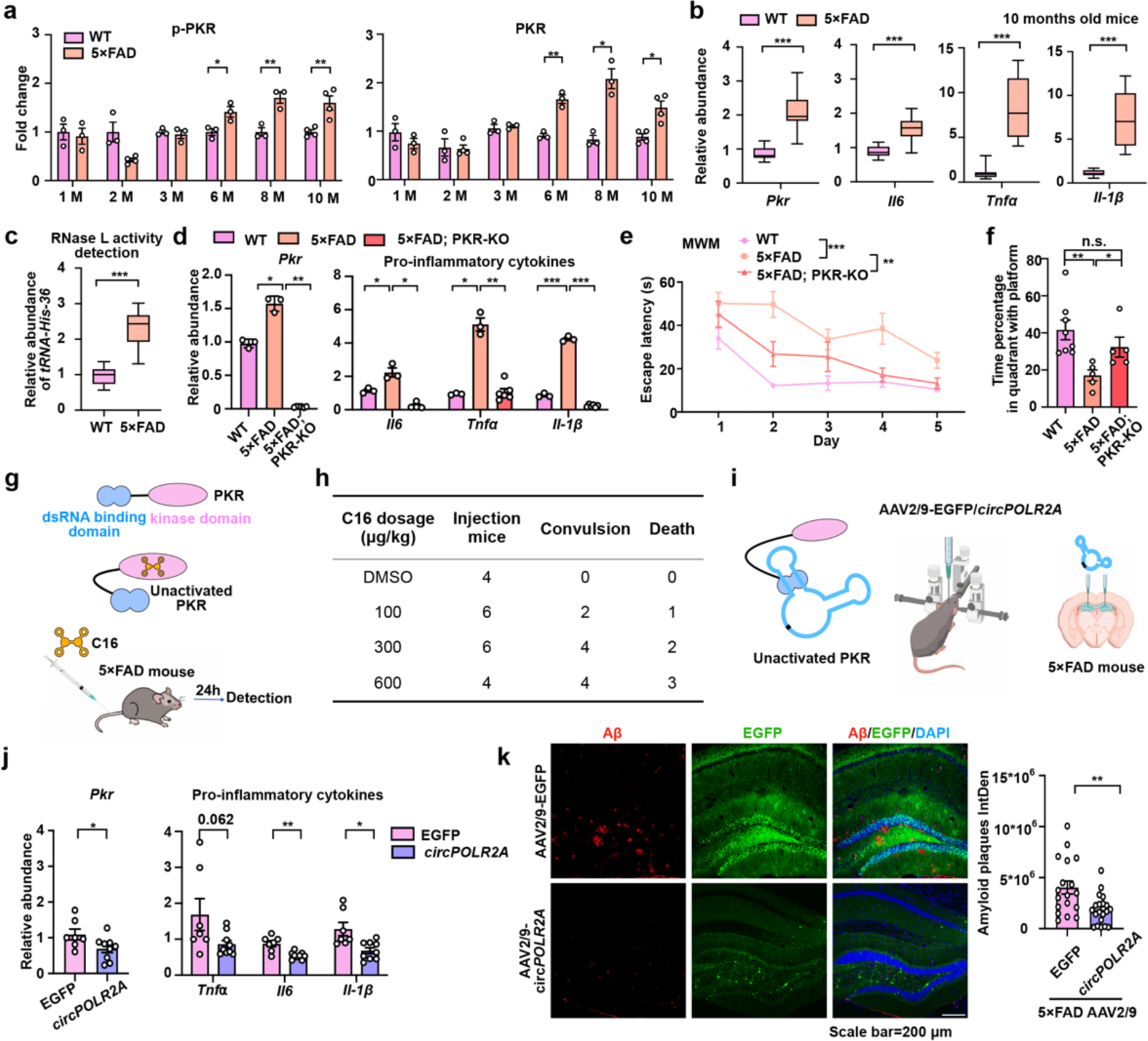
Ds-cRNAs alleviate neuroinflammation by targeting PKR over-activation in 5×FAD mice. **a**, PKR expression and phosphorylation were elevated with AD progression. Levels of PKR and p-PKR were comparable in the hippocampus of 1-month, 2-months, and 3-months-old wild-type (WT) and 5×FAD mice, but elevated in the hippocampus of 6-months, 8-months, and 10-months-old 5ξFAD mice compared to WT mice (n=3 biological repeats). Relative expression of PKR and p-PKR were measured by Western blot (WB). **b**, Expression of *Pkr, Il6*, *Tnfα* and *Il-1β* was higher in hippocampus of 10-month-old 5×FAD mice, compared to WT controls, measured by RT-qPCR. **c**, RNase L activity detection revealed higher activation in 10-month-old 5×FAD mice compared to WT mice, as measured by RT-qPCR. **d**, Compared to 5×FAD mice, the expression of *Pkr, Il6*, *Tnfα* and *Il-1β* in 5×FAD; PKR-KO mice was reduced to a similar level as in WT mouse, measured by RT-qPCR, n=3 biological repeats. **e**, 5×FAD; PKR-KO mice showed better spatial learning ability compared with 5×FAD mice. WT, 5×FAD and 5×FAD; PKR-KO mice underwent Morris Water Maze (MWM). During the hidden platform training sessions of MWM, 5×FAD exhibited a significant learning deficit compared with WT mice. n = 8 (WT), n = 5 (5×FAD), n = 5 (5×FAD; PKR-KO), **p < 0.01, ***p < 0.001. Mean values ± SEMs are shown. Data were analyzed by two-way ANOVA. **f**, Memory deficits were rescued by PKR depletion in 5×FAD mice. Probe trials were conducted to assess the retention of spatial memory. n = 8 (WT), n = 5 (5×FAD), n = 5 (5×FAD; PKR-KO). **g**, 8-month-old 5×FAD mice were injected by tail intravenous with DMSO or C16 at different doses and sacrificed for measurement after 24h. Left, an illustration showed C16 blocked the ATP binding site of the kinase domain for PKR phosphorylation^40^. **h**, Statistics of clonic convulsion and survivals upon DMSO or C16 intravenous injection with different doses in mice. **i**, Left, an illustration showed ds-cRNA interacting with dsRNA binding motifs (dsRBMs) of PKR to prevent its activation without interfering with the kinase domain of PKR. Right, a schematic showed 8-month-old 5×FAD mice received bilateral intrahippocampal injection of AAV2/9-*circPOLR2A* or the control AAV2/9-EGFP. **j**, AAV2/9-*circPOLR2A* delivery to neurons led to reduced expression of *Pkr* and proinflammatory cytokines *Tnfα*, *Il6* and *Il-1β* in the hippocampus of 5ξFAD mice, measured by RT-qPCR, n=7∼9 biological repeats. **k**, AAV2/9-*circPOLR2A* delivery led to decrease Aβ plaques. Left, Aβ plaques in the hippocampus were determined by staining Aβ (anti-Aβ, red), EGFP (green) and DAPI (blue). Right, quantification of Aβ plaques from 18 matching brain sections of 3 mice. **a**, **b**, **c**, **d**, **f**, **j** and **k**: n.s., *p < 0.05, **p < 0.01, ***p < 0.001. two-tailed Student’s t test, Error bars represent mean ± SEM.

Intriguingly, the augmented PKR activation reciprocally accelerated the accumulation of neurotoxic Aβ aggregation, as demonstrated by Encephalomyocarditis virus (EMCV) infection promoting Aβ accumulation in the hippocampus of 3-month-old 5×FAD mice (Extended Data Fig.1d-f).

Next, we generated *5*×*FAD^+/-^*; *Pkr*^-/-^ (refer as 5×FAD; PKR-KO) mice by crossing 5×FAD mice with PKR knockout mice (*Pkr^-/-^*)^19^. The PKR knockout was achieved by deleting 121 bp in exon 3 of *Pkr* using CRISPR/Cas9, resulting in a frameshift mutation and subsequent depletion of PKR^20^. Given that phenotypes were obvious at 8 months old in AD mice (Fig. 1a and Extended Data Fig. 1a-c), most analyses were performed at this age. Western blot analyses of brain samples confirmed the absence of PKR expression in 5×FAD; PKR-KO mice (Extended Data Fig.2a). To assess the mitigation of neuroinflammation in the hippocampus of 5×FAD; PKR-KO mice, RT-qPCR analysis was performed. *Pkr* and several pro-inflammatory cytokines showed a dramatic decrease in 5×FAD; PKR-KO mice compared to their 5ξFAD littermates (Fig. 1d), which even lower than WT mice, suggesting the augmented neuroinflammation during mice aging (Fig. 1d and Extended Data Fig.1b, f). Of note, 5ξFAD; PKR-KO mice displayed improved spatial learning and memory abilities compared to 5×FAD mice, as revealed by the Morris water maze (MWM) (Fig. 1e-f), with no difference in swimming speed or defect in locomotor activity (Extended Data Fig.2b-d). These findings confirmed PKR as an effective target for AD treatment.

Next, we investigated the impact of PKR inhibitors on 5×FAD mice, firstly focusing on small chemical molecules, such as compound #16 (C16), a widely used inhibitor that binds to the kinase domain of PKR (Fig. 1g). Intravenous injections of varying C16 doses (100, 300, and 600 μg/kg) were administered to 5×FAD mice. However, this approach resulted in adverse effects, including convulsions and mortality within 10 minutes post-injection, with severity increasing at higher doses (Fig.1h). Analysis of *Pkr* and pro-inflammatory genes expression in the hippocampus after 24 hours revealed a significant decrease in the 300 μg/kg group (the 600 μg/kg group was excluded owing to only one survival) (Extended Data Fig.3a). Although C16 demonstrated potential in alleviating neuroinflammation in 5×FAD mice, we observed no effects on gliosis, neuroprotection, or Aβ loads following 24 hours of C16 treatment (Extended Data Fig.3b-c). Moreover, severe side effects limited its long-term administration, rendering it unsuitable for AD therapy.

In contrast to most molecular chemicals, ds-cRNAs efficiently suppress PKR activation by preventing its dimerization^20^ (Fig.1i) with no observable toxicity in animals^19^. Thus, we examined the *in vivo* potential of ds-cRNAs to alleviate AD phenotypes by adeno-associated virus (AAV) delivery. AAV-EGFP injection into hippocampus did not trigger detectable inflammatory responses 48 hrs post injection compared to PBS controls (Extended Data Fig.3d-e). AAV2/9-mediated delivery resulted in approximately 50% uptake into neurons, less than 20% into microglia, and the remaining 30% in other cell types (Extended Data Fig.3f). As the human ds-cRNA, *circPOLR2A(9,10)*^12^, was not expressed in mice, we administrated this ds-cRNA in 5ξFAD mice to eliminate background contamination. Sequence of *circPOLR2A(9,10)* was introduced into the circRNA expression plasmid with AAV inverted terminal repeat (ITR) sequence for packaging (Extended Data Fig.3g). The insert sequence is driven by a CMV promoter, allowing a non-cell type-specific expression. It is worthwhile noting that this cassette simultaneously produces ds-cRNA and EGFP by back-splicing and splicing, respectively^21, 22^, thus EGFP was used as a marker to indicate the successful ds-cRNA production (Fig. 1k).

We firstly delivered *circPOLR2A(9,10)* into bilateral hippocampus by intrahippocampal injection with AAV2/9 (Fig. 1i). Upon intrahippocampal injection with AAV2/9, Northern blot confirmed successful delivery of *circPOLR2A(9,10)* into the hippocampus (Extended Data Fig.3h). As expect, we observed only a mild reduction of *PKR* mRNA, but a dramatic decrease of PKR activation, a remarkable reduction of pro-inflammatory cytokines and Aβ plaques in the hippocampus upon *circPOLR2A(9,10)* addition (Fig.1j-k and Extended Data Fig.3i-j). All AAV2/9-*circPOLR2A(9,10)* treated mice via intrahippocampal injection were viable (Extended Data Fig.4b). These findings suggest the potential application of ds-cRNAs in targeting p-PKR-mediated neuroinflammation and Aβ plaque formation, showing a safety advantage over the examined small molecule inhibitor C16.

We then utilized tail intravenous injection for efficient whole-brain delivery via AAV-PHP.eB, which can pass the blood-brain barrier^23^. Another human ds-cRNA, *circCAMSAP1(2,3)*, was also introduced to eliminate potential sequence-specific effects of *circPOLR2A(9,10)* in mice. Northern blot confirmed positive ds-cRNA expression (Extended Data Fig.4a). Brain samples from the injected mice revealed a slight decrease in PKR expression, accompanied with significantly reduced PKR activation and pro-inflammatory cytokines expression in 5×FAD mice administered with ds-cRNAs post four weeks, compared to EGFP controls (Fig.2a-b). Furthermore, the ds-cRNA-treated 5×FAD mice exhibited notably decreased Aβ plaques in the hippocampus compared to EGFP-injected controls (Fig.2c). Delivery of the two different ds-cRNAs, respectively, both resulted in dramatically reduced proliferation (Fig.2d) of microglia. Importantly, all mice remained viable and exhibited normal behavior throughout the experimental period (Extended Data Fig.4b).

**Fig. 2.**
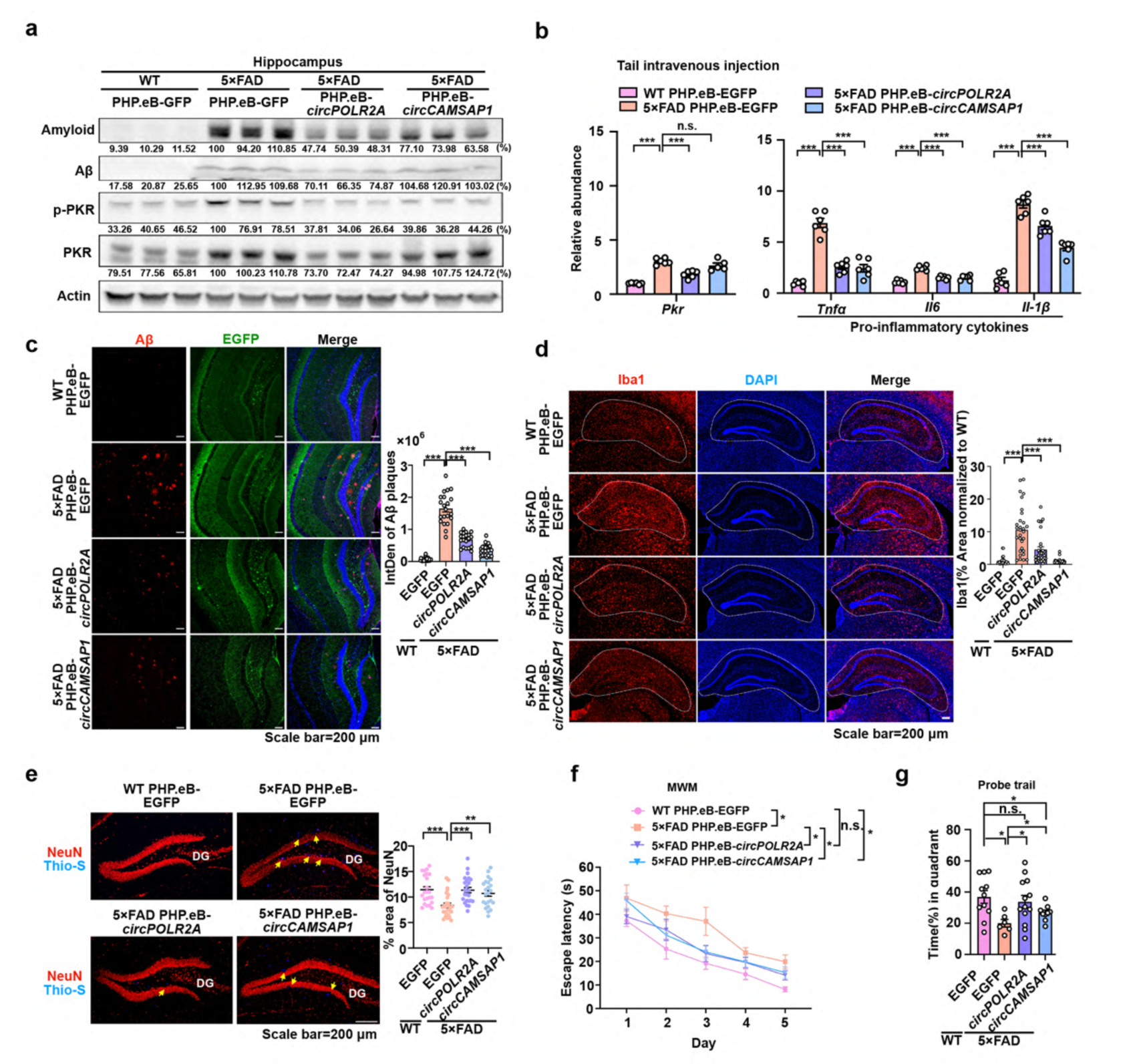
Ds-cRNAs ameliorate AD-relevant phenotypes. **a**, Delivery of ds-cRNAs by AAV-PHP.eB-*circPOLR2A* and AAV-PHP.eB-*circCAMSAP1*, individually, dampened PKR over-activation and Aβ expression in the hippocampus of 5×FAD mice. n=3 biological repeats. Relative expressions of PKR, p-PKR and Aβ were measured by WB. **b**, Delivery of ds-cRNAs (*circPOLR2A* and *circCAMSAP1*) by AAV-PHP.eB, individually, attenuated inflammatory responses in the hippocampus of 5×FAD mice, as revealed by RT-qPCR. **c**, AAV delivery of ds-cRNAs led to alleviated Aβ plaque formation in the hippocampus of 5×FAD mice. Left, representative images of Aβ plaque composition labeled with Η31L21 (anti-Aβ, red), DAPI (blue) and EGFP (green). The presence of EGFP indicated the expression of respect ds-cRNAs given the co-occurance of splicing and back-splicing simultaneously (See Extended Data Fig.3f and Li et al., 2017^22^). Right, quantification represented Aβ plaque level in the hippocampus from 6∼10 matching brain sections per mouse, n=3∼4. **d**, AAV delivery of ds-cRNAs alleviated gliosis in 5ξFAD mice. Left, representative immunofluorescence images of hippocampal sections from 8–9-month-old WT mice injected with PHP.eB-EGFP, and 5× FAD mice injected with PHP.eB-EGFP, PHP.eB-*circPOLR2A* or PHP.eB-*circCAMSAP1* stained with antibodies to Iba1 (red) and DAPI (blue). Right, the relative percentage of the area of sections taken from the hippocampus covered by Iba1 staining compared with WT group. Mean values ± SEMs are shown (n = 21 to 32 slices from 3-4 mice per group). **e**, Ds-cRNAs addition prevented neuron loss (yellow arrows) in the hippocampus of 5xFAD mouse. Left, a representative image of neuron (anti-NeuN, red) and Aβ (anti-Aβ, blue) in the DG region in hippocampal section injected with AAV-PHP.eB EGFP or ds-cRNAs. Yellow arrows identified the neuronal loss with Aβ plaque aggregation. Right, Statistics of the percentage of NeuN^+^ area relative to the total hippocampal area, n = 23∼27. **f**, 5×FAD mice injected with AAV PHP.eB-*circPOLR2A* or AAV PHP.eB-*circCAMSAP1* displayed increased spatial learning ability than those injected with AAV PHP.eB-EGFP in 8-9 months-old. Mice were conducted for behavioral test one month after injection using the hidden platform training sessions of MWM. n = 11 (WT PHP.eB-EGFP), n = 6 (5×FAD PHP.eB-EGFP), n = 13 (5×FAD PHP.eB-*circPOLR2A*), n = 9 (5×FAD PHP.eB-*circCAMSAP1*), *p < 0.05. Mean values ± SEMs are shown. Data were analyzed by two-way ANOVA. **g**, Memory retention deficits were rescued by overexpression with ds-cRNA in 5×FAD mice. Probe trials were conducted to assess the retention of spatial memory. n = 11 (WT PHP.eB-EGFP), n = 6 (5×FAD PHP.eB-EGFP), n = 13 (5×FAD PHP.eB-*circPOLR2A*), n = 9 (5×FAD PHP.eB-*circCAMSAP1*). **b**, **c**, **d**, **e**, and **g**: n.s., *p < 0.05, **p < 0.01, ***p < 0.001. two-tailed Student’s t test, Error bars represent mean ± SEM.

Due to the propensity of Aβ^24^ and overactivated microglia^25^ to impair neuronal function, we hypothesized that ds-cRNAs addition could enhance neuroprotection. As expected, ds-cRNA-treated 5×FAD groups displayed approximately a 1.5-fold increase in neurons in the dentate gyrus (DG) region of hippocampus compared to 5×FAD controls, comparable to WT controls at the age of 8 to 9 months (Fig.2e). Subsequently, we assessed whether the AAV-ds-cRNA treatments could ameliorate spatial learning and memory defects in the 5×FAD mice. The ds-cRNA-administered 5×FAD mice showed a remarkably improvement in spatial learning and memory abilities compared to EGFP-injected 5×FAD controls during the MWM test (Fig.2f-g). Of note, these behavioral rescues were not due to the altered swimming speed or the changes of locomotor activity (Extended Data Fig.5). Moreover, treatment with ds-cRNA at both early stage (3 months old 5×FAD mice) and late stage (10 months old 5×FAD mice) of AD exhibited improvements in AD phenotypes (Extended Data Fig.6). In addition, the frontotemporal dementia mouse model, PS19, characterized by super-phosphorylation of Tau with mutant hTAU^26^, also showed significant rescue in AD pathologic phenotypes and improvement in spatial learning and memory with ds-cRNAs addition (Extended Data Fig.7). Of note, activated PKR activation was shown to promote Tau tangle^27^. Collectively, these results indicated that delivery of ds-cRNAs as PKR inhibitors *in vivo* efficiently dampen PKR activation, alleviating AD phenotypes and neuroinflammation, improving the spatial learning and memory abilities in two different models.

To further illustrate the potential therapeutic application of ds-cRNAs in AD, we conducted immunofluorescent analysis of p-PKR to identify cell types undergoing PKR activation. We found nearly 6.6% of p-PKR was presented in microglia, while the majority (87%) of p-PKR signal was detected in neurons (Fig.3a). This distribution aligns with the known PKR expression pattern in the human brain^28^ (www.alzdata.org) and is corresponded with our earlier observation that AAV primarily targets neurons (Extended Data Fig.3f). On the other hand, the portion of microglia in the normal adult mouse brain only 3.7% (Fig.3b). 6.6% p-PKR located in microglia suggested a large portion of microglia accompanying with activated PKR. Moreover, microglial (CD11b^+^) are the major contributors to neuroinflammation^29, 30^, secreting pro-inflammatory cytokines dozens of times higher than CD11b^-^ samples in 5ξFAD mice (Extended Data Fig.8a). We further characterized microglia during ds-cRNA-mediated AD therapy.

**Fig. 3.**
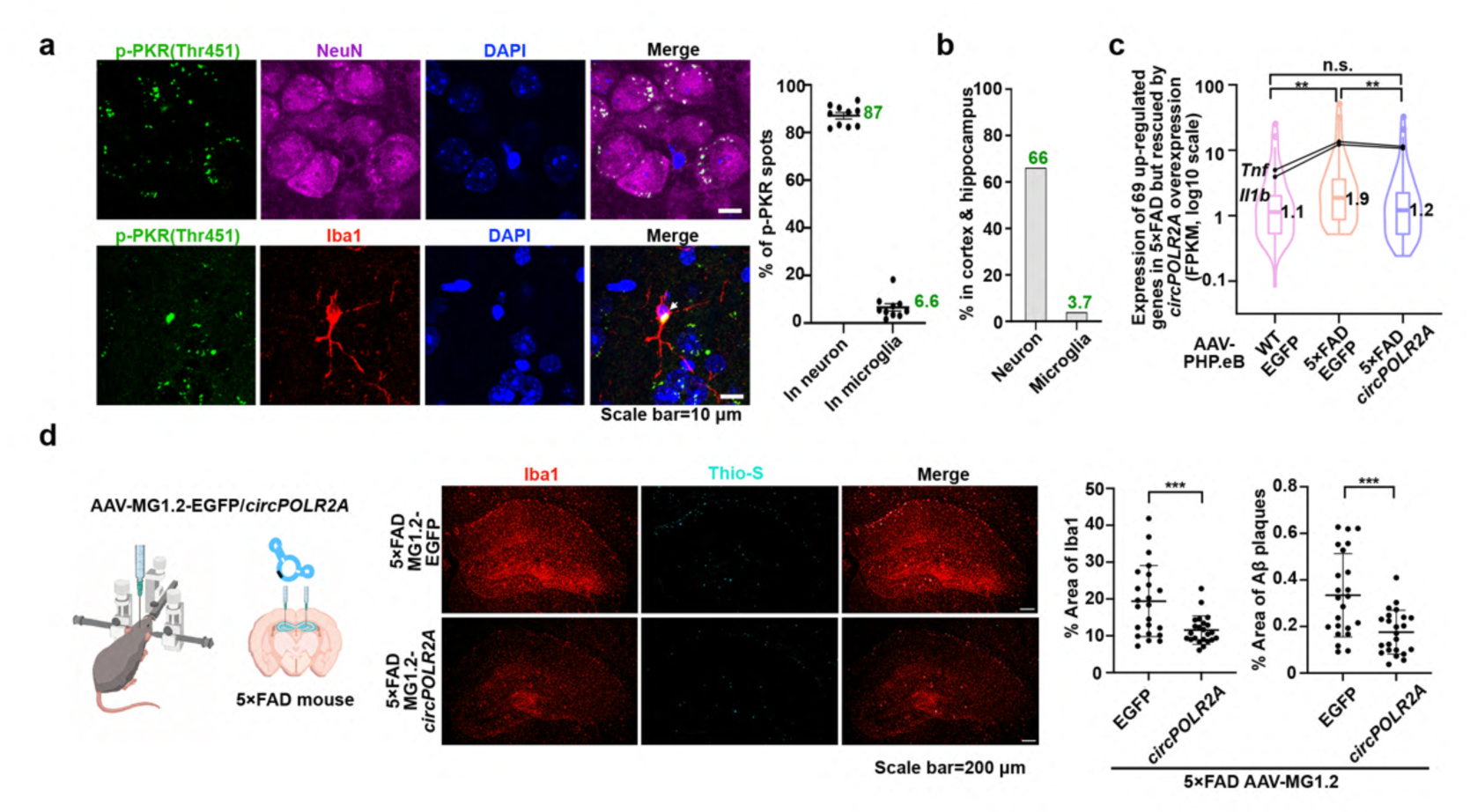
Ds-cRNAs rescue AD phenotypes by targeting neurons and microglia. **a**, p-PKR puncta mainly accumulated in neurons in 8-month-old 5×FAD mice hippocampus. Images were acquired using confocal microscopy of p-PKR (green), co-stained with neurons (NeuN positive; violet), microglia (Iba1 positive; red) and counterstained with DAPI in the hippocampus of 5×FAD mice. Scale bar: 20μm. **b**, The proportion of neuron and microglia in mouse cortex and hippocampus. (Data were from the mouse whole-brain transcriptomic cell type atlas in Allen Brain Cell Atlas database). **c**, Expression distribution of 69 up-regulated genes in microglia of 5× FAD mice, which were rescued by *circPOLR2A* delivery. The medians, IQRs, and 1.5×IQRs and outliners exceeding 1.5×IQRs are shown. Wilcoxon rank-sum test. n.s., **p < 0.01. **d**, Microglia-specific *circPOLR2A* delivery reduced microglia proliferation and decreased Aβ plaques in hippocampus. Left, a schematic of 8-month-old 5×FAD mice injected with AAV-MG1.2-*circPOLR2A* or the control AAV-MG1.2-EGFP into bilateral hippocampus by intrahippocampal injection. Middle, representative immunofluorescence images of hippocampal sections from 8–9-month-old 5xFAD mice injected with MG1.2-EGFP or MG1.2-*circPOLR2A* stained with Iba1 (red) and Aβ (Thio-S, cyan). Right, the statistics of the percentage of Iba1 and Aβ positive area relative to the total hippocampal area. Mean values ± SEMs are shown (n = 23 slices from 3 mice per group). Data were analyzed by two-tailed Student’s t test. ***p < 0.001.

Firstly, microglia from 5×FAD mice or WT controls 4 weeks post AAV-PHP.eB injection were isolated respectively, followed by total RNA collection and ribosomal RNA (rRNA)-depleted RNA-seq (Extended Data Fig.8b). In general, 930 upregulated and 1,791 downregulated genes were presented in microglia from the 5×FAD mice compared to those from WT ones (Extended Data Fig.8c). Gene ontology (GO) revealed 36% (334/930) of all upregulated genes involved in immune response pathways (Extended Data Fig.8d), including *Tnf* and *Il-1b* (Fig.3c). AAV-*circPLOR2A(9,10)* delivery rescued 7.42% (69/930) up-regulated genes comparing to 5ξFAD controls (Extended Data Fig.8c,e), of which 23.2% (16/69) up-regulated genes are related to immune responses (Fig.3c, Extended Data Fig.8e). Further analyses showed genes involved in chemical synaptic transmission and cation channel function (34.8%, 24/69) were also rescued to at least 1.5-fold decrease in expression (Extended Data Fig.8e), consistent with the behavior test results (Fig.2f-g).

Notably, delivery of *circPOLR2A(9,10)* specifically into microglia in bilateral hippocampus of 5×FAD mice via AAV-MG1.2^31^ also led to fewer microglial cells, one month later, (Fig.3d) compared to EGFP controls. Consistently, mice receiving *circPOLR2A(9,10)* exhibited decreased Aβ plaques labeled by Thio-S in the hippocampus compared to EGFP controls (Fig.3d). Although mechanism of the Aβ decrease via reduced p-PKR awaits additional investigation, our data suggests that targeting neuroinflammation and mitigating the overactivation of proliferated microglia, which might enhance the production and propagation of Aβ in late-stage AD^32, 33^, representing a promising therapeutic strategy for AD. Collectively, this microglia-specific ds-cRNA addition and RNA-seq data corroborate our characterization of ds-cRNA as an efficient PKR inhibitor^19^ capable of ameliorating AD-relevant phenotypes (Fig.1, 2).

In sum, we propose AAV-delivered ds-cRNAs as a novel strategy for AD therapy by targeting excessive PKR activation, dampening neuroinflammation and AD progression (Extended Data Fig.8f). It is important to note potential limitations, including the reliance on the two distinct mouse models that each partially mimics AD phenotypes and the challenge of long-term AAV treatment due to AAV capsid immunogenicity. Future therapeutic exploration and expanded indications may benefit from delivery of immunogenicity-free ds-cRNAs^19^ via brain-specific lipid nanoparticles (LNPs).

## Acknowledgement

We thank Boxun Lu for discussion and reading the manuscript, Minmin Luo and Rui Lin for providing packaging plasmids of AAV-MG1.2, Naihe Jing for gifting 5×FAD mice, Shiqing Cai and Yuejun Chen for gifting PS19 mice, Zhuohao He for gifting antibody anti-PHF1. This work was supported by the National Key R&D Program of China (2021YFA1300501), the Strategic Priority Research Program of the Chinese Academy of Science (XDB0570000), Science and Technology Commission of Shanghai Municipality (23DX1900100; 23DX1900101) to L.-L.C. This work has been supported by the New Cornerstone Science Foundation through the New Cornerstone Investigator Program and the XPLORER PRIZE.

## Author information

L.-L.C. supervised and conceived the project. X.F. and B.-W.J. designed and performed experiments. S.-N.Z. preformed computational analyses, supervised by L.Y. and L.-L.C., J. W. generated the NGS library. B.-Q.Z. and M.-Y. W. helped with mice experiments. L.-L.C., X.F., B.-W.J., S.-N.Z., C.-X.L., and H.W. wrote the paper with input from all authors. All authors read and approved the manuscript.

## Ethics declarations

L.-L.C., X.F., B.-W. J., and C.-X.L. are named as inventors on patents related to circular RNA aptamers held by CAS Center for Excellence in Molecular Cell Science (CEMCS).

## Methods

### Experimental animals

5×FAD mice on a C57BL/6J background were provided as gifts by Naihe Jing at Guangzhou institute of biomedicine and health, Chinese Academy of Sciences. PS19 mice on a C57BL/6J background were provided as gifts by Yuejun Chen and Shiqing Cai at Center for Excellence in Brain Science and Intelligence Technology, Chinese Academy of Sciences. PKR-KO mice were generated in our lab. Wild-type C57BL/6J mice were purchased from Shanghai Laboratory Animal Center, Chinese Academy of Sciences (Shanghai, China). All animal experiments were approved by the Committee of Use of Laboratory in the Center for Excellence in Molecular Cell Science, Chinese Academy of Sciences. Mice were raised together with littermates in pathogen-free environment, and their health status was routinely checked. No more than 5 mice were housed in one cage. The mice were housed in standard cages with free access to food and water. Mice were maintained in a 12-h light/dark cycle at 22–26°C. Experiments were conducted during the light phase of the cycle. Animals were randomly assigned to different groups. Male mice of 8-month-old were used for all injections and behavior tests. To genotype the mice, genomic fragments were amplified from tail DNAs using designed specific primers (Supplementary Table S2).

### Human cell lines

The human cell line, A549, was purchased from the American Type Culture Collection (ATCC; http://www.atcc.org) and was cultured in F-12K (Gibco) supplemented with 10% Fetal Bovine Serum (Gibco) and 1% GlutaMax (Gibco).

### Bacterial strains

E. coli expression strain BL21 [Transetta (DE3) chemically competent cell] were procured from Transgen Biotech (Cat# CD801) and were grown in LB culture at 37°C.

### Virus strains

The overexpressed virus AAV2/9 and AAV-MG1.2 used in this study was packaged and produced by BrainVT A (Wuhan) Co., Ltd, AAV-PHP.eB was packaged and produced by Guangzhou PackGene Biotech Co., Ltd and BrainVT A (Wuhan) Co., LTD. The packaging plasmids of AAV-MG1.2 were provided as gifts by Minmin Luo and Rui Lin in National Institute of Biological Sciences (NIBS), Beijing, China.

### Plasmid constructions

When constructing the AAV overexpression plasmid, DNA fragment of 1/2 EGFP-complementary sequence-POLR2A-complementary sequence-1/2EGFP was obtained from the pZW1-FCS-circRNA^21^ plasmid by PCR, and then was inserted into the pAAV-MCS plasmid vector by homologous recombination method. Plasmid of *circCAMSAP1(2,3)* was constructed in the same way. After that, transformation and plasmid extraction were performed. Primers for plasmid constructions were listed in Table S2. All constructs were confirmed by Sanger sequencing.

### Cell culture

A549 and N2A cell lines were maintained using standard protocols from ATCC.

### RNA isolation, RT-qPCR

Total RNA from tissues was extracted using TRIzol reagent (Invitrogen) and subjected to cDNA synthesis using PrimeScript^TM^ RT Master Mix (TaKaRa). qPCR was performed using QuantStudio6 Flex System (Thermo Fisher) and THUNDERBIRD® SYBR® qPCR Mix (TOYOBO). GAPDH mRNA was examined as an internal control for normalization. Expression of each examined gene was determined from three independent experiments. Primers for RT-qPCR were listed in Supplementary Table S2.

### Northern blotting (NB)

NB was performed according to the manufacturer’s protocol (DIG Northern Starter Kit, Roche). Digoxigenin (Dig)-labeled antisense riboprobes were produced using using T7 RNA polymerase by in vitro transcription with RiboMAX Large Scale RNA Production Systems (Promega). In brief, 10 μg total RNAs or 5 ng in vitro synthesized linear or circular RNAs was resolved on denaturing urea polyacrylamide gel, transferred to nylon membrane (Roche) and UV-crosslinked using standard manufacturer’s protocol. Membrane was then hybridized with specific Dig-labeled riboRNA probes. NB probes were listed in Supplementary Table S2.

### Library preparation and deep sequencing

For RNA-seq samples from mouse microglia, total RNAs were processed with RiboMinus kit (Human/ Mouse Module, Invitrogen) to deplete most ribosomal RNAs (ribo-RNA). The ribo-RNA libraries for sequencing were prepared with illumina TruSeq® Stranded Total RNA Library Prep Gold (#20020598) kit. All libraries were sequenced with NovaSeq 6000 system at Sequanta Technologies. FastQC was applied for raw read qualities assessment.

### Western blotting (WB)

Tissues were treated with SDS-PAGE loading buffer at 95°C for 10 minutes. The samples were then separated on an SDS-PAGE gel, transferred onto a PVDF membrane, and probed with specific antibodies. The protein bands of interest were then visualized using enhanced chemiluminescence (Tanon), and the intensity of the bands was quantified using Fiji software.

### Tissue processing and sample preparation

For immunofluorescence assay, mice were euthanized and transcranial perfused with 25 mL of 1×PBS, 25 mL of 4% paraformaldehyde in 37°C and 25 mL of 4% paraformaldehyde in 4°C. Brains were dissected out, then drop-fixed in 4% paraformaldehyde over night at 4°C. Drop-fixed samples were transferred to 25% sucrose for 24 h and 30% sucrose for 48 h, then mounted and frozen in Epredia™ Neg-50™ Frozen Section Medium (Fisher scientific). These brains were then sectioned at 25 μm in thickness using a cryostat (Leica) and stored in PBS at 4 °C for downstream staining and imaging. For RNA and protein extraction, hippocampus and cortex were dissociated, flash-frozen and stored at −80°C. Hippocampal tissues were mechanically homogenized in 300 μL of RIPA buffer (50 mM Tris-HCl pH 7.6, 15 mM NaCl, 1% NaNoc, 0.1% SDS, 1% NP40, 1 mM EDTA, 1mM EGTA, 1 M NaF, 0.1 M Na_3_VO_4_, 100 mM PMSF, 0.1 mM DTT) containing EASYpack Protease Inhibitor Cocktail (Roche). Following homogenization, the samples were spun down at 15000 rpm for 10 min and 100 μL of the supernatant were isolated for western blotting. Remaining supernatant and pellet were diluted in 1 mL Trizol for RNA extraction.

### Immunofluorescence staining of brain samples

Floating brains sectioned stored in PBS were blocked with 10% donkey serum, 0.125% Gelatin, 0.05% Tween 20 in PBS for 2 h at room temperature before applying the primary antibody master mix diluted in antibody diluent solution (1% BSA, 0.03% Triton X-100 in PBS) overnight at 4°C. Samples were stained with anti-Ab (H31L21, Invitrogen, 1:1000, 6Ε10, BioLegend, 1:1000) to label plaques. To study microglial morphology and numbers, sections were stained with Iba1 (019-19741, Wako, 1:1000) and NeuN (MAB377, Millipore Sigma, 1:500). To characterize EMCV infection, we labeled with anti-phospho-tau (AT8, Thermo Fisher, 1:500) and anti-dsRNA (J2, SCICONS, 1:200). Sections were then washed 3 times for 10 min at room temperature in PBS, then incubated in matched donkey Alexa Fluor 488, 594, 647 anti-rabbit, -goat, and -mouse (Jackson, 1:1000 dilution) at 37°C for 30 min. Samples were washed again 3 times for 10 min at room temperature and incubated with DAPI (1:1000) for 10 min at room temperature, or stained for plaques with Thioflavin S (Sigma-Aldrich, 0.1% in 80% ethanol) for 8 min followed by three 2-min washes with 50% ethanol at room temperature. All tissue sections were then transferred to wells containing PBS before being mounted to glass slides with VECTASHIELD® antifade mounting medium (Vector Lab) and coverslips. Mounted slides were stored at 4°C until imaged using LAS AF software (Leica Microsystems) on a Leica TCS SP8 confocal microscope. Quantification of images were performed using Fiji software.

### MACS isolation of microglia

Mice were transcranial perfused using ice cold Hank’s balanced salt solution (HBSS) (Gibco) and the brains were quickly dissected, then hippocampal and cortical tissues were placed in HBSS on ice. The tissues were cut into small pieces using a sterile scalpel and centrifuge at 300× g for 2 min at room temperature and aspirate the supernatant carefully. Small pieces of tissues were digested with 10 U/mL papain (Worthington Bio Corp) and 0.1 mg/mL DNase I (Sigma) for 18 min at 37 °C. After removing digestive solution, the digestion was terminated by adding 1 mL of 10% FBS DMEM culture medium, and the tissues were gently pipetted 15-20 times with a pipette. Centrifuge at 300× g for 5 min to collect the sample at the bottom of the tube, wash with MACS buffer (0.5% BSA, 1mM EDTA in PBS) and filter through a 70-μm strainer (CSS013070, Biofil).

After the cell isolations, the cell pellets were subjected to a 15-minute staining procedure in the dark using Anti-CD11b Magnetic Microbeads (130–093-634, Miltenyi Biotec) on ice. The cells were then washed with MACS buffer and centrifuged at 400 ×g for 10 minutes. Subsequently, the cell pellet was resuspended in 500 μl of MACS buffer. The autoMACS Pro Separator (130-092-545, Miltenyi Biotec) was primed, and both the sample and collection tubes were placed in a cold MACS Chill 5 Rack (130-092-951, Miltenyi Biotec). Both positive and negative fractions were collected using the separator. After the separation, the positive fraction containing the cells of interest was preserved for further RNA-seq analysis.

### Intrahippocampal injection

5×FAD, PS19 and WT mice were anesthetized before receiving a bilateral hippocampal injection of 2 μL of ΑΑV2/9, AAV-MG1.2 or EMCV into the right and left hemisphere of the hippocampus (at ±1.5 mm lateral, −2.0 mm posterior, and −2.2 mm ventral relative to the intersection of the coronal and sagittal suture (bregma) at a rate of 200 nL/min) using a stereotaxic frame. For AAV expression assay, 1 month post injection, mice were euthanized and transcranial perfused before preparing brains for immunofluorescent staining to evaluate Aβ clearance in the hippocampus. Images were analyzed using Fiji software. For EMCV infection assay, 24 h post injection mice were sacrificed for protein and RNA extraction.

### Behavior tests

All behavior experiments were performed between 8 AM and 8 PM in a blinded fashion. All mice were 9∼10 months old at the time of the assay. Mice were transported from their home vivarium room to the behavior core and allowed one hour to habituate before beginning each test.

### Open field test

The open field test is a widely used method for assessing exploratory behavior and general activity in animals. To create a conducive testing environment, the room was dark and sound insulated. The tracking instrument utilized infrared lasers and sensors to identify and monitor the central body point of each mouse accurately. The mice were gently introduced into a white plastic open-field arena measuring 40cm × 40cm × 40cm. A video camera positioned directly above the arena tracked the movement of each mouse, and the data was recorded for a 10-minute duration on a computer using EthoVision XT software. This software was used to measure and analyze two key aspects: the total distance traveled by the mice and the time spent in the center of the chamber compared to the time spent near the edges. Anxiety-like behavior was assessed based on the mice’s preference for spending more time at the edges of the box and less time in the center. Longer durations at the edges were interpreted as potential signs of anxiety. After each trial, the arena was cleaned using 70% ethanol to maintain cleanliness and prevent potential confounding effects between different test subjects.

### Rotarod

Rotarod testing is a common experimental method used to evaluate motor coordination, balance, and motor learning in laboratory animals. In this study, WT and AD mice treated with AAV were subjected to an accelerating rotarod test. During the test, the rotarod apparatus (YLS-4C) was set to gradually increase the rotation speed from 0 to 40 rpm over a period of 90s. Subsequently, the speed was maintained at a constant level. The time elapsed before the mouse fell off the rotating rod was recorded, with the maximum observation time set at 5 minutes. Each animal underwent a session consisting of 3 trials per day, with at least a 30-minute interval between each trial. The data from the 3 trials were averaged to provide a more reliable assessment. The latency of the mice to fall from the rod was used as an index to evaluate their motor coordination. This rotarod testing protocol allows us to measure and compare motor performance between the WT and AD mice, providing valuable insights into potential motor impairments associated with Alzheimer’s disease and the effects of AAV treatment on motor functions.

### Morris water maze(MWM)

To assess spatial learning and memory, we conducted the Morris Water Maze (MWM) test following established protocols^34^. The water tank was partitioned into four quadrants, each featuring a distinct geometric symbol (pentagram, square, triangle, and circle) affixed to its respective wall, serving as external cues for spatial orientation. The water temperature was carefully maintained at 22 ± 1°C, and we introduced food-grade titanium dioxide particles into the water to facilitate animal tracking. The experimental apparatus was placed in a controlled environment free from noise and intense light sources.

Before commencing the experiment, mice were acclimated to the testing environment for a duration of 1 hour. Subsequently, each mouse underwent a training regimen spanning five consecutive days, encompassing four trials per day, with a minimum one-hour interval between trials. The mice were introduced into the water maze, initially positioned with their heads facing the pool’s inner wall, and the location of the hidden platform was randomly assigned to one of the four quadrants. We recorded the time it took for the animals to locate the platform during each training trial. If a mouse exceeded a latency of 60 seconds in locating the platform during a training session, they were gently guided to the platform and allowed to remain there for 30 seconds. If the mice find the platform within 60 seconds, then stay on the platform for 5-10 seconds.

This training protocol was repeated for a total of 5 days to ensure that mice had ample opportunities to learn the platform’s location. Twenty-four hours after training, the platform was removed and the 30-s probe test commenced. During this test, mice were placed into the water maze, facing the quadrant opposite to the target quadrant. We recorded the amount of time the mice spent in the target quadrant as an indicator of their spatial memory performance.

### Ribo-RNA-seq processing and analysis

Pair-end RNA-seq reads were firstly trimmed by Trimmomatics^35^ (v0.32, parameters: PE-threads 5-phred33 ILLUMINACLIP:TruSeq3PE-2.fa:2:30:10 LEADING:3 TRAILING:3 SLIDINGWINDOW:4:15 MINLEN:36) to remove adaptors and low-quality bases. Next, the reads were mapped to the mm10 rDNA sequences for rRNAs (5S, 5.8S,18S, 28S) and pre-rRNAs (45S) to remove reads from ribosomal RNA by Bowtie2^36^ (version: 2.3.5, parameters: -p 10 --score-min L,-16,0 --rfg 0,7 --rdg 0,7 --mp 7,7) and the rest paired reads were then mapped to the mouse mm10 genome reference with HISAT2^37^ (version 2.1.0, parameters: --no-softclip --score-min L,-16,0 --mp 7,7 --rfg 0,7 --rdg 0,7 --max-seeds 20 --dta -k 1). Fragment counts of genes were obtained using featureCounts featureCounts^38^ (v1.5.1, parameters: -s 2 -p --fraction -O -T 16 -t exon -g gene_id) based on gencode vM25 gene annotation and normalized to FPKM (fragments per kilobase of gene per million fragments mapped)^37^. Differentially expressed genes were determined with a threshold of fold change ≥ 1.5 between two conditions. The gene ontology enrichment analyse were performed by clusterProfiler^39^ (version 3.14.3, parameters: pvalueCutoff = 0.05, qvalueCutoff = 0.2).

### Reporting summary

Further information on research design is available in the Nature Portfolio Reporting Summary linked to this article.

## Data Availability

Mouse microglia RNA-seq datasets are available at Gene Expression Omnibus (GSE249440). The mm10 reference genome and gencode vM25 annotation GTF file were download from GENCODE database (GRCm38.p6). All original unprocessed data related to this paper were uploaded to Mendeley Data.

## Code availability

This paper does not report original code.

## Supplementary information

**Supplementary Table S1.** Gene expression (FPKM) for microglia RNA-seq.

**Supplementary Table S2.** List of construct, probe and primer sequences.

---

**Extended Data Fig. 1.**
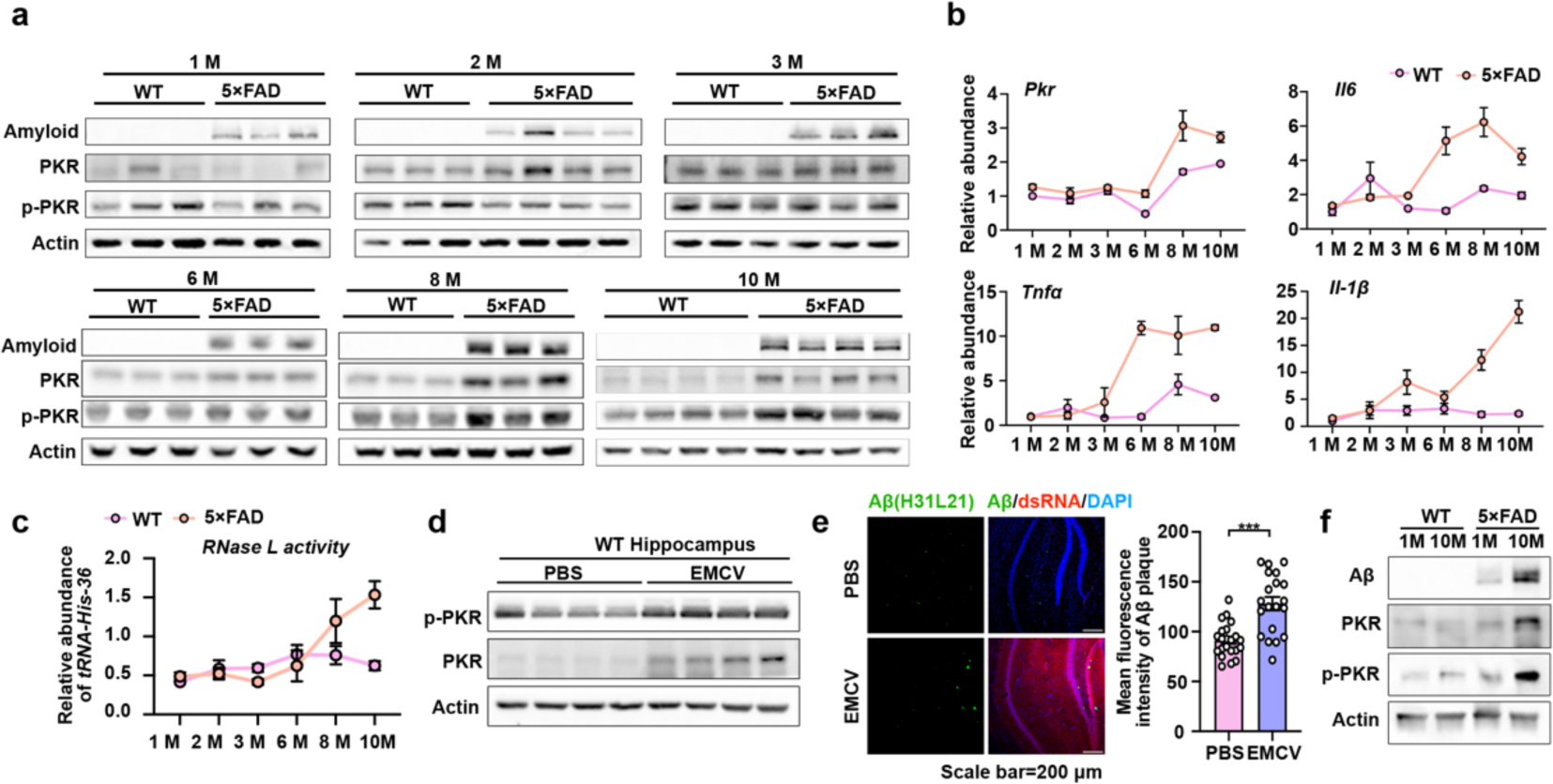
Elevated PKR with AD progression exacerbated neuroinflammation and Aβ aggregation. **a**, PKR and p-PKR levels were comparable in hippocampus of 1 month, 2 months and 3 months old wild type (WT) and 5×FAD mice, but increased in the hippocampus of 6 months, 8 months and 10 months old 5×FAD mice, compared with WT mice. n=3 biological repeats. Relative expression of PKR and p-PKR is measured by WB. **b**, Expression of *Pkr*, *Il6*, *Tnfα* and *Il-1β* was comparable in 1 month, 2 months and 3 months old mice and became up-regulated in 5×FAD mouse hippocampus at 6-month-, 8-month- and 10-month-old. n=3 biological repeats. Relative expression of *Pkr*, *Il6*, *Tnfα* and *Il-1β* were measured by RT-qPCR. **c**, RNase L activity remained comparable in 1-month-old, 2-month-old, and 3-month-old mice, but became up-regulated in the hippocampus of 5×FAD mice at 6 months, 8 months, and 10 months of age, as measured by RT-qPCR. n=3 biological repeats. **d**, Increased phosphorylation of PKR was triggered by EMCV infection. n=4 biological repeats. Relative expression of PKR and p-PKR were measured by WB. **e**, EMCV infection triggered Aβ plaques formation in the hippocampus of 5×FAD mice. Left, representative images of Aβ plaques formation (labeled by H31L21 in green) in 3-month-old 5×FAD mouse hippocampus upon EMCV (labeled by J2 anti-dsRNA in red) stimulation. Right, quantification of Aβ in hippocampus from 6∼8 matching brain sections per mouse. **f**, PKR and p-PKR levels were increased in both WT and 5×FAD mice as they aged or with AD progression. Detected by WB with 1-month- and 10-month-old WT and 5×FAD mice.

**Extended Data Fig. 2.**
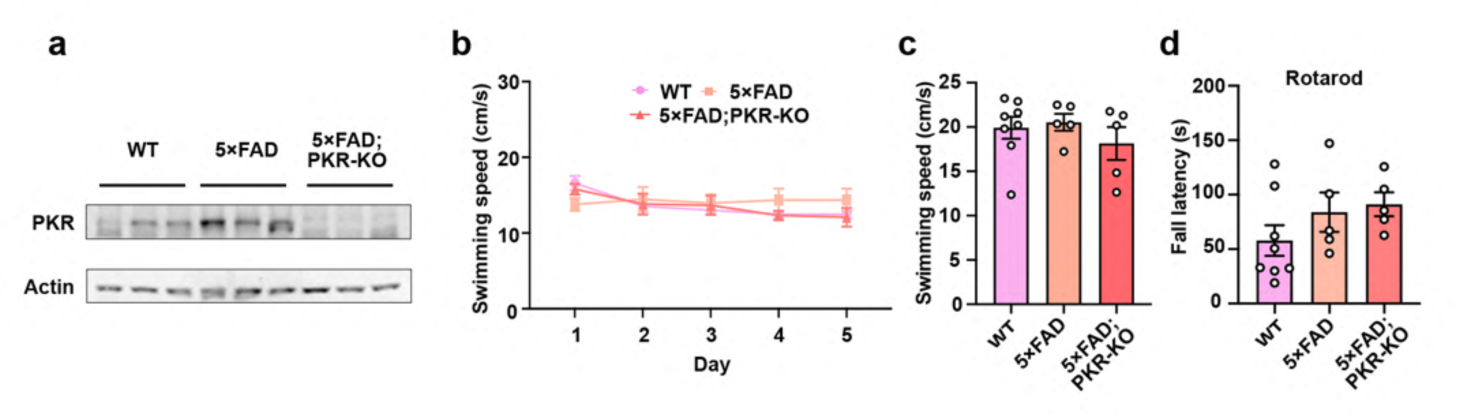
5×FAD; PKR-KO mice show comparable motor abilities in behavior tests. **a**, PKR knockout was confirmed by WB in 5×FAD; PKR-KO mice. **b-c**, Swim speed in MWM showed no significant difference. Mean values ± SEMs are shown. **d**, Motor function test on rotarod showed intact motor balance activity of all mice. **b**-**d**: n = 8 (WT), n = 5 (5× FAD), n = 5 (5× FAD; PKR-KO). Mean values ± SEMs are shown. Data were analyzed by two-tailed Student’s t test.

**Extended Data Fig. 3.**
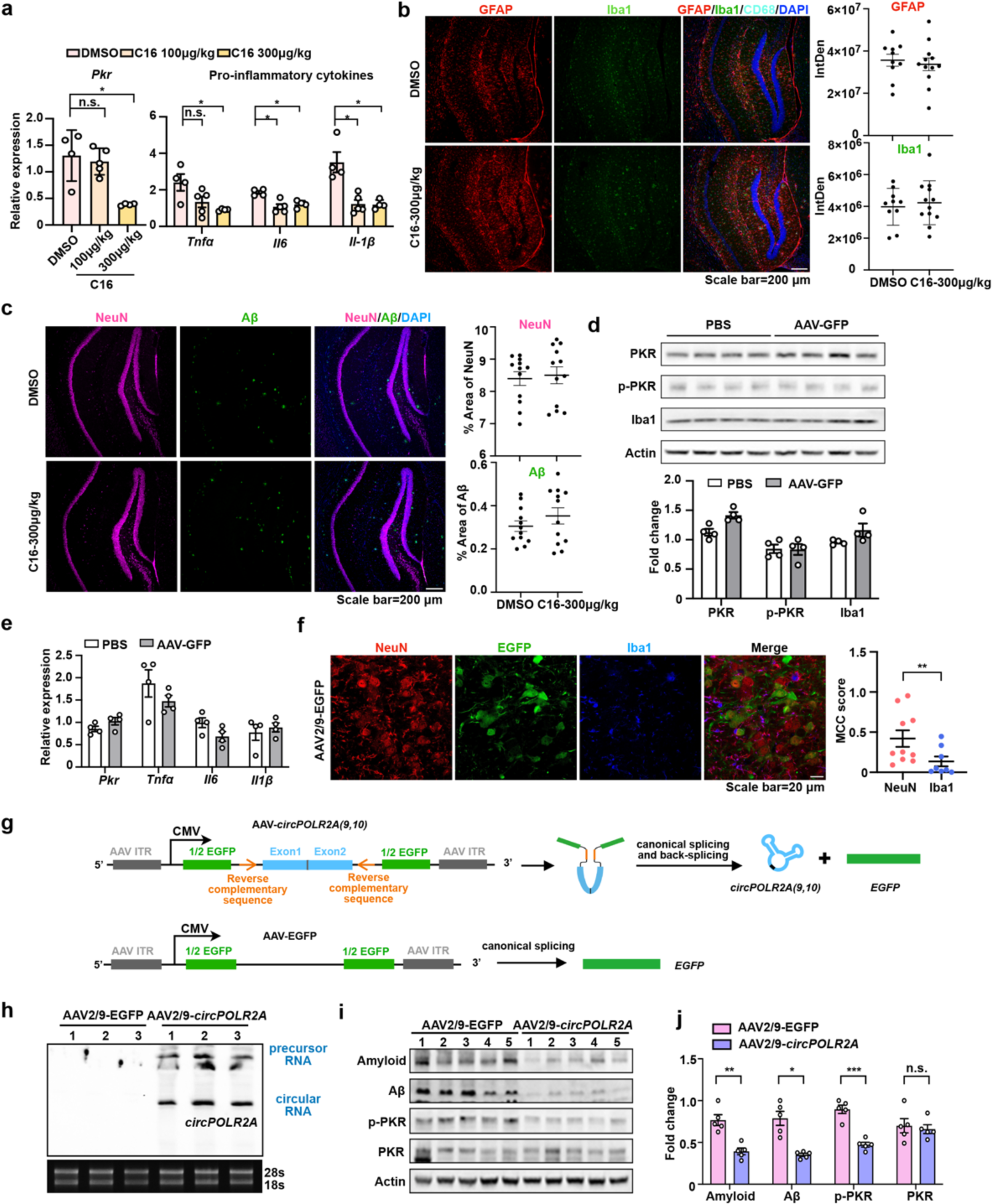
Ds-cRNA outcompetes C16 to alleviate molecular symptoms in 5×FAD mice. **a**, C16 in 300 μg/kg prevented *Pkr* and pro-inflammatory cytokines *Il6*, *Tnfα* and *Il-1β* expression in the hippocampus of 5×FAD mice, as detected by RT-qPCR. **b**, IF showed that treatment with C16 (300 μg/kg) did not rescue gliosis in 5×FAD mice. Left showed representative pictures. Right, quantification of microglia (Iba1in green) and astrocyte (GFAP in red) in hippocampus. **c**, IF showed that treatment with C16 (300 μg/kg) did not improve neuroprotection or Aβ clearance. Left showed representative pictures. Right, quantification of neuron (NeuN in purple) and Aβ (6E10 in green) in hippocampus. **d,** AAV infection did not activate PKR or stimulate microglia proliferation. Above, PKR phosphorylation (p-PKR-T446, p-PKR for simplicity), PKR expression and microglia marker Iba1 expression were assayed by WB. Below, quantification of PKR, p-PKR and Iba1 expression in WB. The levels were normalized by Actin expression, respectively. **e**, AAV infection did not cause detectable inflammatory responses in hippocampus of WT mice, as shown by RT-qPCR. **f**, IF staining showed that AAV-EGFP mainly infects to neurons(NeuN marked in red) and parts of microglia (Iba1 marked in blue). Right, colocalization analysis of EGFP and NeuN or Iba1 measured via Manders’ colocalization coefficients (MCC) from 6 brain sections per mouse by IamgeJ. **g**, The illustration showed the vectors used for AAV packaging to express ds-cRNA or EGFP. Top, an illustration showed the vector of *circPOLR2A* containing exons of *POLR2A*, 1/2 *EGFP* fragments, reverse complementary sequences facilitating back splicing, and AAV inverted terminal repeats (ITR) for AAV packaging, producing *circPOLR2A* and *egfp* mRNA. Bottom, an illustration showed the vector of EGFP as control, producing *egfp* mRNA only. **h**, *CircPOLR2A* expressed in hippocampus of mice with AAV intrahippocampal injection, as revealed by Northern Blot (NB) on denaturing PAGE. **i-j**, AAV2/9-*circPOLR2A* delivery attenuated excessive PKR phosphorylation and reduced Aβ plaques in the hippocampus of 5×FAD mice. **a**, **b**, **c**, **d**, **e**, **f** and **j**: n.s., *p < 0.05, **p < 0.01, ***p < 0.001. Two-tailed Student’s t test, Error bars represent mean ± SEM.

**Extended Data Fig. 4.**
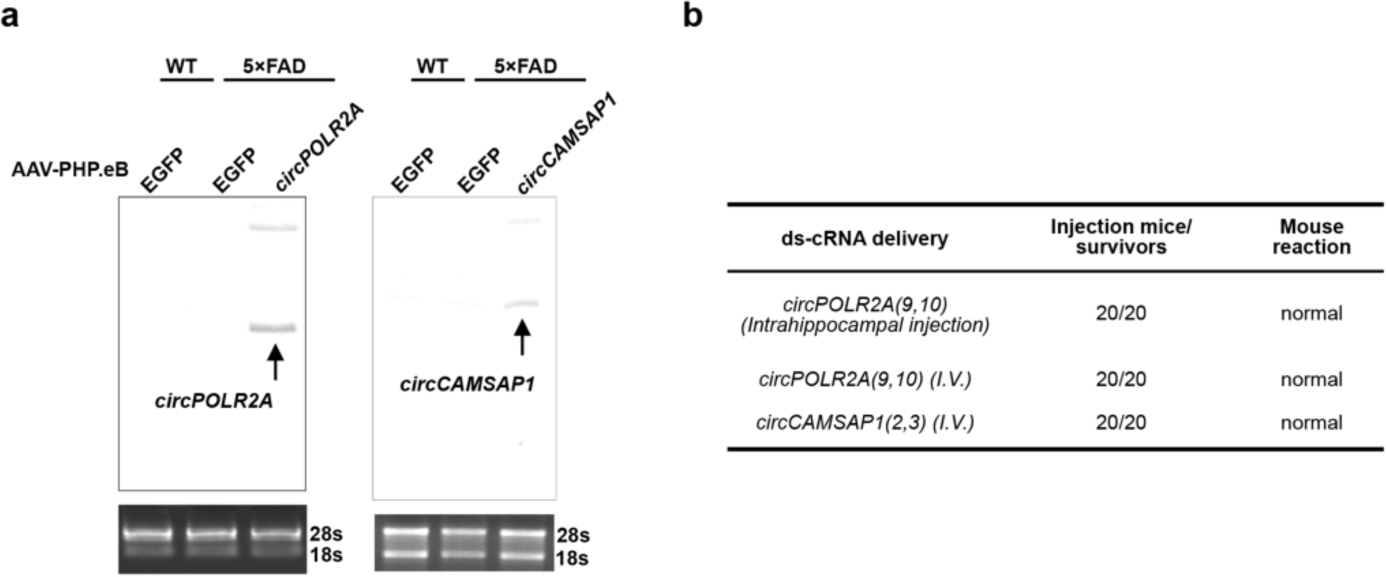
Delivery of ds-cRNAs into the hippocampus by intravenous injection in 5×FAD mice. **a**, *CircPOLR2A* and *circCAMSAP1* were expressed in the hippocampus by intravenous injection (*I.V.*) of AAV-PHP.eB, as confirmed by NB on denaturing PAGE. **b**, Summary of mice situation upon ds-cRNA delivery with AAV utilizing different injection methods.

**Extended Data Fig. 5.**
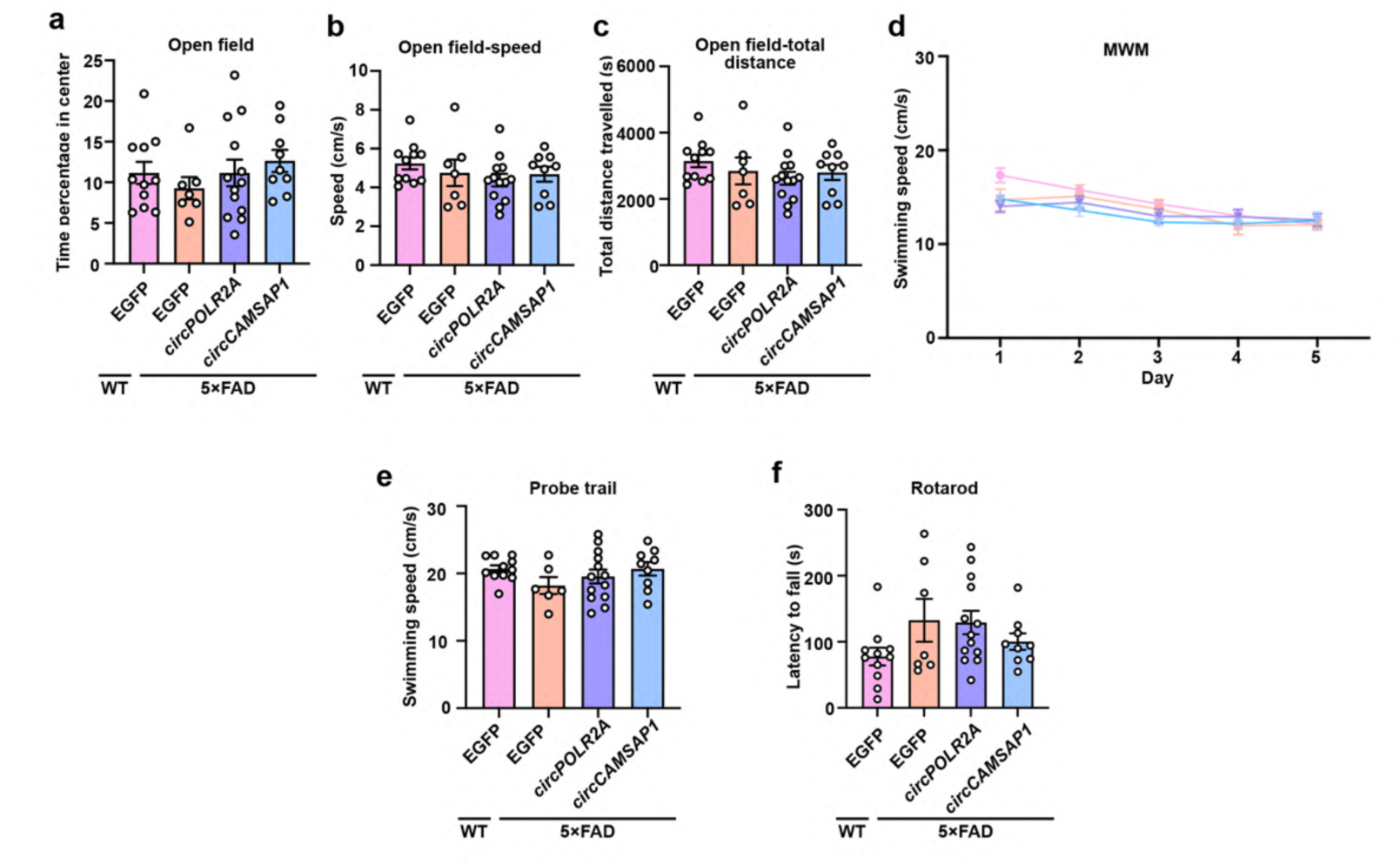
5×FAD mice with ds-cRNAs delivered by AAV intravenous injection show no difference in anxiety and motor ability. **a-c**, Open field test. Time percentage spent in center (a), moving speed (b) and total distance of movement (c) in an open field for 10 min. n = 11 (WT PHP.eB-EGFP), n = 6 (5×FAD PHP.eB-EGFP), n = 13 (5×FAD PHP.eB-*circPOLR2A*), n = 9 (5×FAD PHP.eB-*circCAMSAP1*). Behavioral mice showed no anxiety. Data were analyzed by two-tailed Student’s t test. **d-e**, Swim speed in the Morris water maze showed no significant difference. Data were analyzed by two-tailed Student’s t test. **f**, Motor function was tested on the rotarod. Data were analyzed by two-tailed Student’s t test. Error bars represent mean ± SEM.

**Extended Data Fig. 6.**
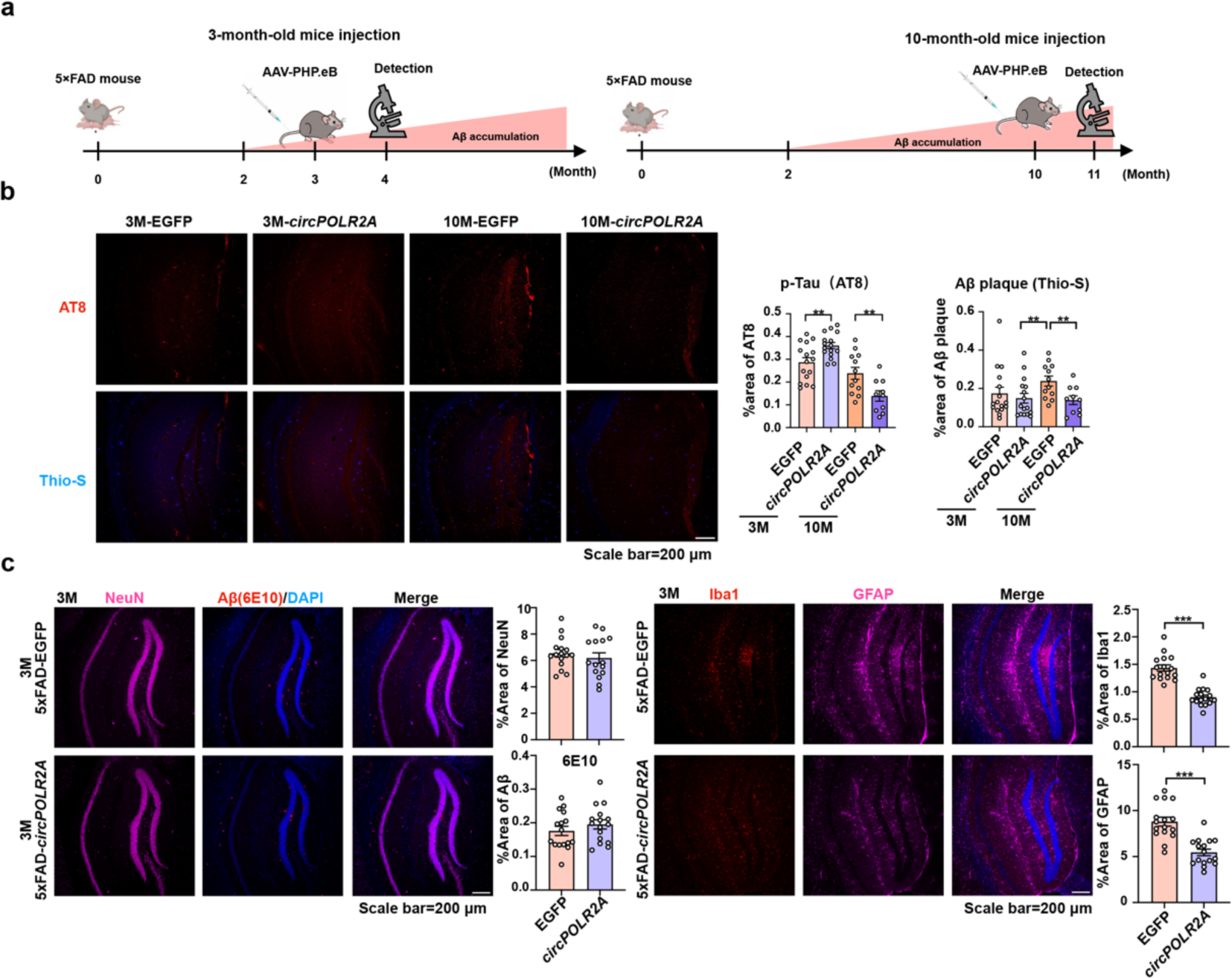
Adding ds-cRNAs at either the early or late stage of AD rescues AD phenotypes in 5×FAD mice. **a**, The diagrams illustrate the different timelines of AAV injection in 5×FAD mice. **b**, AAV delivery of ds-cRNAs alleviated Aβ and p-Tau aggregation in 10-month-old 5×FAD mice, but not in 3-month-old mice. On the left are representative immunofluorescence images of hippocampal sections from 3-month-old and 10-month-old 5×FAD mice injected with PHP.eB-EGFP and PHP.eB-circPOLR2A, stained with AT8 (red) and Thio-S (blue). On the right are the relative percentages of the area of sections taken from the hippocampus covered by AT8 and Thio-S staining compared with EGFP controls. Data were analyzed using a two-tailed Student’s t-test. **c**, In 3-month-old 5×FAD mice, the addition of ds-cRNAs decreased gliosis but did not affect neuron death or Aβ clearance. On the left are representative immunofluorescence images, and on the right are the relative percentages of the area of sections taken from the hippocampus covered by NeuN, Aβ, Iba1, and GFAP. Data were analyzed by two-tailed Student’s t test.

**Extended Data Fig. 7.**
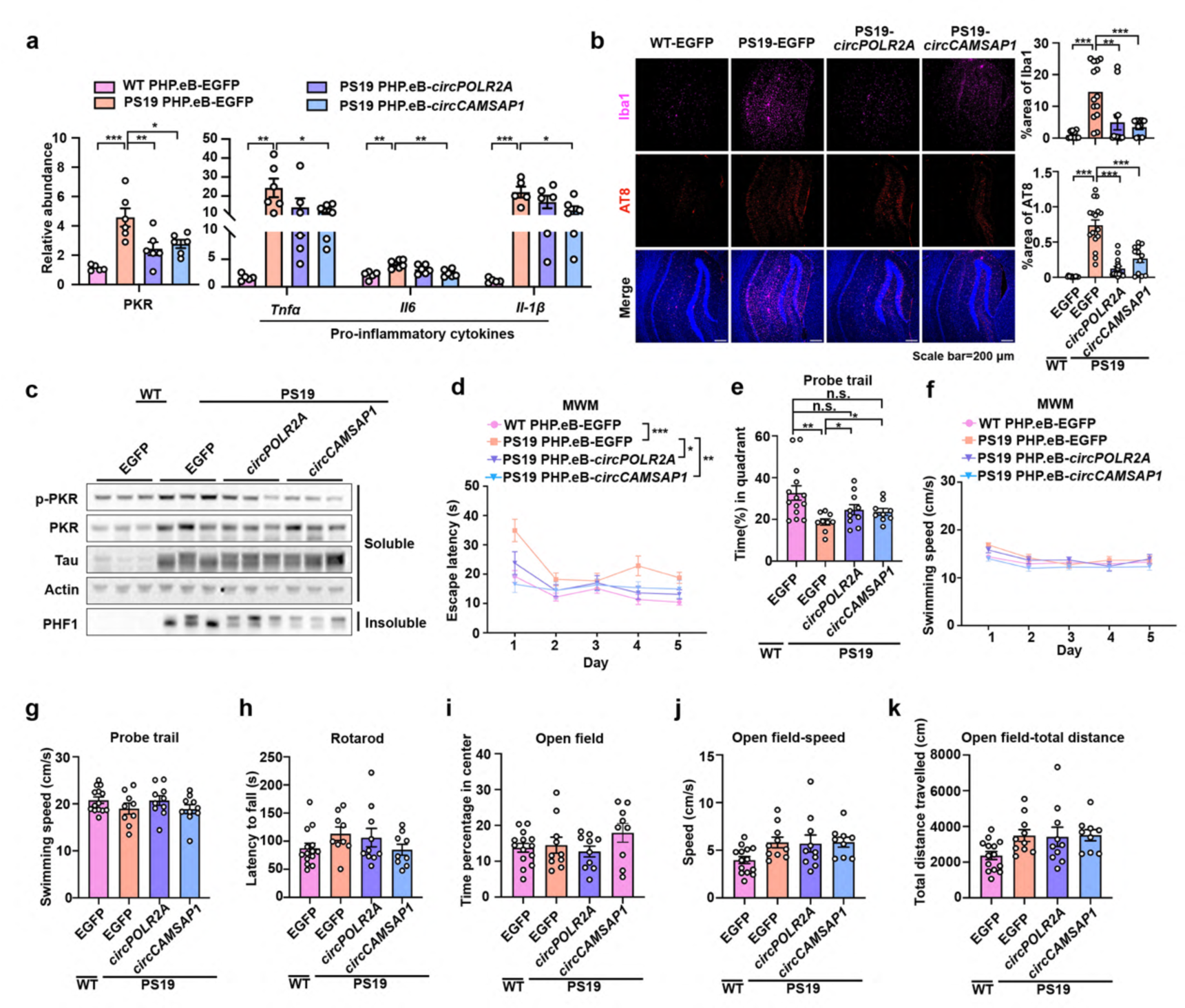
Ds-cRNAs addition by AAV rescues AD phenotypes and memory deficits in PS19 mice. **a**, Delivery of ds-cRNAs (*circPOLR2A* and *circCAMSAP1*) by AAV-PHP.eB, individually, dampened inflammatory responses in the hippocampus of PS19 mice, as revealed by RT-qPCR. Data were analyzed by two-tailed Student’s t test. **b**, AAV delivery of ds-cRNAs alleviated gliosis and p-Tau aggregation in PS19 mice. Left, representative immunofluorescence images of hippocampal sections from 8–9-month-old WT mice injected with PHP.eB-EGFP, and PS19 mice injected with PHP.eB-EGFP, PHP.eB-*circPOLR2A* or PHP.eB-*circCAMSAP1* were stained with Iba1 (violet), p-Tau (AT8, red) and DAPI (blue). Right, the relative percentage of the area of sections taken from the hippocampus covered by Iba1 and AT8 staining compared with the WT group (n = 8 slices from 3-4 mice per group). Data were analyzed by two-tailed Student’s t test. **c**, Delivery of ds-cRNAs by AAV-PHP.eB-*circPOLR2A* and AAV-PHP.eB-*circCAMSAP1*, individually, dampened PKR extra-activation and p-Tau (PHF1) expression in the hippocampus of PS19 mice. **d**, PS19 mice injected with AAV PHP.eB-*circPOLR2A* or AAV PHP.eB-*circCAMSAP1* displayed increased ability in spatial learning compared to those injected with control AAV. PHP.eB-EGFP in 8∼9-month-old WT/PS19 mice through intravenous injection. Behavioral tests were conducted three months later using the hidden platform training sessions of MWM. n = 14 (WT PHP.eB-EGFP), n = 9 (PS19 PHP.eB-EGFP), n = 10 (PS19 PHP.eB-*circPOLR2A*), n = 9 (PS19 PHP.eB-*circCAMSAP1*). Data were analyzed by two-way ANOVA. **e**, Probe trials were conducted to assess the retention of spatial memory. Memory retention deficits were rescued by overexpression with ds-cRNA in PS19 mice. Data were analyzed by one-tailed Student’s t test. **f-g**, Swim speed in the Morris water maze had no significant difference. **h**, Motor function was tested on the rotarod. Data were analyzed by two-tailed Student’s t test. **i-k**, Open field test. Time percentage spent in center (i), moving speed (j) and total distance of movement (k) in an open field for 10 min. Behavioral mice showed no anxiety. Data were analyzed by two-tailed Student’s t test. **a**, **b** and **d**-**k**: n.s., *p < 0.05, **p < 0.01, ***p < 0.001. Error bars represent mean ± SEM.

**Extended Data Fig. 8.**
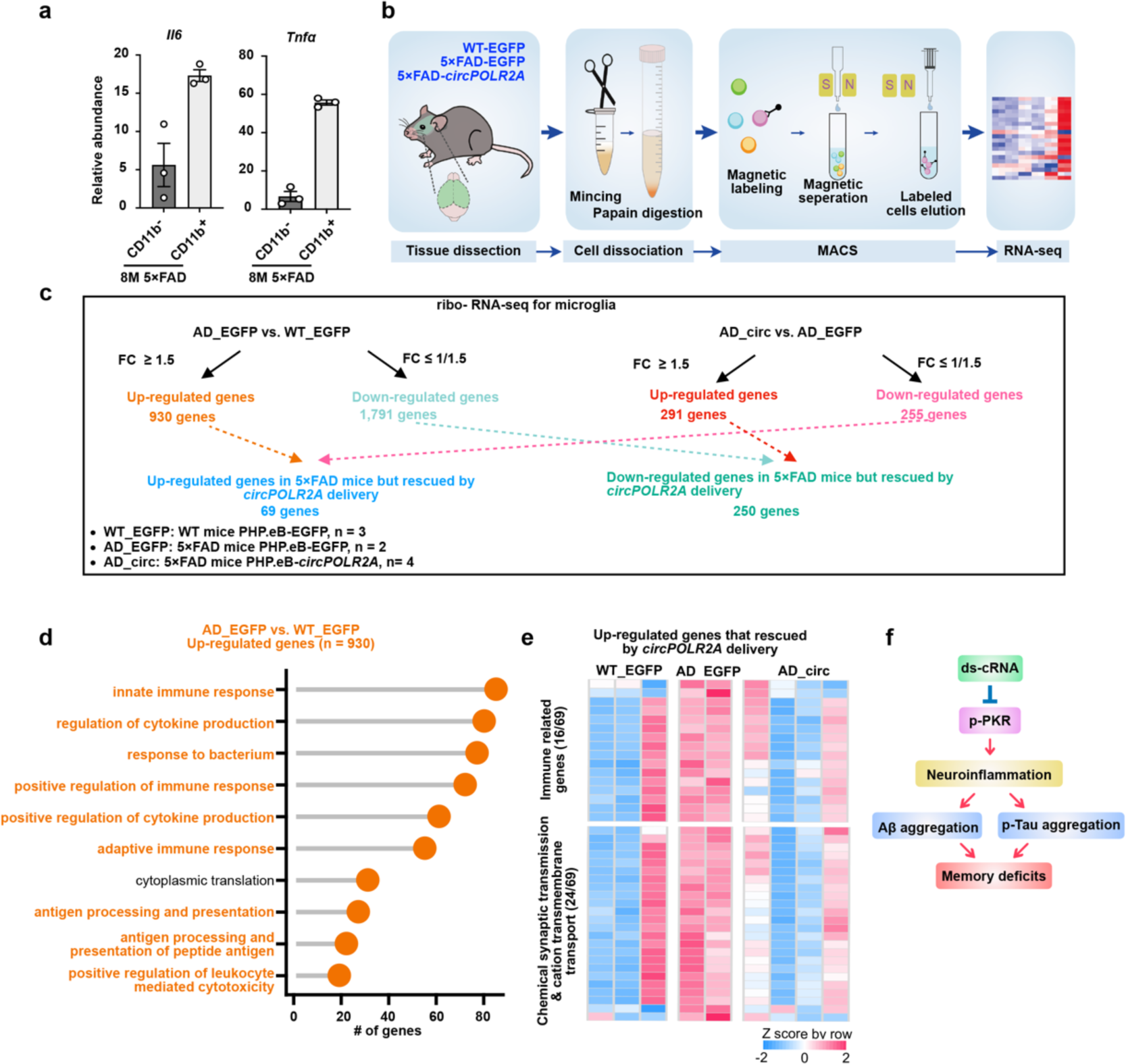
AAV-mediated ds-cRNA delivery alleviates microglia inflammation. **a**, Microglial cells (CD11b^+^) in the brain of 5×FAD mice expressed over ten times higher levels of *Il6* and *Tnfα* compared to other cells (CD11b^-^), as detected by RT-qPCR. **b**, Schematic illustrated the process of microglia isolation and RNA preparation for RNA-seq. **c**, Pipeline for detecting dysregulated genes in microglia of 5×FAD mice, which were rescued by *circPOLR2A* delivery. **d**, GO enrichment analysis for the 930 up-regulated genes in microglia of 5×FAD mice revealed significant enrichment primarily in immune related pathways. **e**, Heatmap illustrated the relative expression levels of 40 out of69 genes rescued by *circPOLR2A* delivery in microglia of 5×FAD. Among them, 16 genes were immune-related, and 24 genes were involved in chemical synaptic transmission and cation transmembrane transport. n = 3 (WT PHP.eB-EGFP mice), n= 2 (5×FAD PHP.eB-EGFP mice), n = 4 (5×FAD PHP.eB-*circPOLR2A* mice) **f**, Schematic showed that ds-cRNA inhibited PKR activation and the p-PKR-mediated inflammation cascade, resulting in reduced aggregation of Aβ and p-Tau^5, 41, 42^, and ultimately alleviated the memory deficits in AD.

